# CRISPR-mediated deletion of the Protein tyrosine phosphatase, non-receptor type 22 (PTPN22) improves human T cell function for adoptive T cell therapy

**DOI:** 10.1101/2020.12.03.410043

**Authors:** Sonja Prade, David Wright, Nicola Logan, Alexandra R. Teagle, Hans Stauss, Rose Zamoyska

**Author notes:** To Whom Correspondence should be addressed: Rose Zamoyska, Institute of Immunology and Infection Research, University of Edinburgh, Ashworth Laboratories Rm 1.13, Charlotte Auerbach Road, Edinburgh EH9 3FL, UK.

## Abstract

Adoptive T cell transfer has improved the treatment of cancer patients. However, treatment of solid tumors is still challenging and new strategies that optimize T cell function and response duration in the tumor could be beneficial additions to cancer therapy. In this study, we deleted the intracellular phosphatase PTPN22 and the endogenous TCR α chain from human PBMC-derived T cells using CRISPR/Cas9 and transduced them with TCRs specific for a defined antigen. Deletion of PTPN22 in human T cells increased the secretion of IFNγ and GM-CSF in multiple donors. The cells retained a polyfunctional cytokine expression after re-stimulation and greater numbers of PTPN22^KO^ T cells expressed inflammatory cytokines compared to unmutated control cells. PTPN22^KO^ T cells seemed to be more polyfunctional at low antigen concentrations. Additionally, we were able to show that that PTPN22^KO^ T cells were more effective in controlling tumor cell growth. This suggests that they might be more functional within the suppressive tumor microenvironment thereby overcoming the limitations of immunotherapy for solid tumors.

## Introduction

Advances in immunotherapy have greatly improved the treatment of cancer in recent years. A variety of cancers have been targeted using T cells engineered to express tumor-specific T cell receptors (TCRs) or chimeric antigen receptors (CARs) and some remarkable responses have been reported especially in hematological malignancies [1–5]. However, effective T cell mediated immunotherapy of solid tumors is beset by a number of challenges. On the one hand, the immunosuppressive tumor microenvironment may interfere with T cell function [6]. Cells in the tumor produce anti-inflammatory cytokines, including IL-10 and TGFβ, that can be highly suppressive to effector T cells. As an example, T cells that were engineered to be refractory to TGFβ signaling showed enhance effector function [7]. Additionally, tumors frequently contain high numbers of regulatory T cells (Tregs) that can suppress effector T cell responses [8]. On the other hand, persistent exposure to the tumor antigen may result in T cell exhaustion [9]. Indeed, TCR-engineered T cells that accumulated in the tumor progressively lost the ability to produce IFNγ and TNFα[10]. In addition, cancers can evade immune surveillance by downregulating the expression of tumor antigens [11, 12]. Thus, tumors frequently fail to stimulate robust T cell responses and promote long-lasting T cell anti-tumor activity [13].

Strategies that promote durable T cell responses which overcome these challenges would be beneficial adjuncts to adoptive cell therapy. Moreover, an understanding is required of what type of effector T cell response is most effective against different types of cancers. It was suggested that higher-affinity TCRs were associated with better anti-tumor responses [14]. Higher-affinity and higher peptide concentrations are associated with T cells that are more polyfunctional, meaning that these cells are able to simultaneously produce multiple cytokines [15] and elicit more effective T cell responses against tumors [16, 17]. However, high-affinity TCRs specific for tumor-associated antigens (TAAs) are rarely present in the human T cell repertoire, because tolerance mechanisms select against them [18]. Moreover, TAAs are generally present at fewer than 50 copies per tumor cell, so their effective concentration is low [19, 20]. Together the combination of low affinity TCRs and scarcity of TAAs make activation of polyfunctional T cells less likely to occur. Therefore, we took an alternative approach to improve the function of human T cells for anti-tumor cell therapy. We engineered T cells that are more responsive to low-affinity antigens, which are present at low concentrations, by removing a negative regulator of T cell activation, the intracellular tyrosine phosphatase, PTPN22. In addition, we tried to achieve fine-tuning of effector functions in order to avoid side effects like cytokine release syndrome [21]. PTPN22 curtails TCR signaling, particularly in response to weak antigens, and polymorphisms in the gene have been identified as risk factors for autoimmunity [22, 23]. Murine CD8 T cells lacking PTPN22 were shown to be resistant to wild-type Treg-mediated suppression [24] and to the suppressive effects of TGFβ [25], characteristics that make them superior to wildtype CD8 T cells in controlling the growth of established tumors [25, 26].

In this study, we transduced primary PBMC-derived human T cells with TCRs specific for a defined antigen and used CRISPR/Cas9 to delete endogenous TCR and PTPN22 genes. PTPN22^KO^ T cells were tested for pro-inflammatory cytokine production and effector function. Deletion of PTPN22 in human T cells increased the secretion of IFNγ and GM-CSF in multiple donors. The cells retained a polyfunctional cytokine expression after re-stimulation and greater numbers of PTPN22^KO^ T cells expressed inflammatory cytokines (IFNγ, GM-CSF, TNFα and IL-2) compared to unmutated control cells. Additionally, we were able to show that PTPN22^KO^ T cells were more effective in controlling tumor growth in a murine transfer model. Therefore, human PTPN22^KO^ T cells behave similarly to mouse PTPN22^KO^ T cells, they are more polyfunctional and are likely to be more resistant to suppressive signals from the tumor microenvironment thereby improving the potential of T cell immunotherapy for the treatment of solid tumors.

## Results

### Multiplex genome editing disrupts PTPN22 and TCR expression in primary human T cells

In order to generate a population of tumor antigen specific CD8 T cells, we took the approach of replacing the endogenous TCR in normal donor peripheral blood T cells with a transduced TCR specific for common tumor antigens in the context of HLA-A2. A common approach to reduce off-target cytotoxicity of the transferred T cells is to knockout the endogenous TCR. To this end, we decided to use a multiplex genome editing approach to delete both PTPN22 and the endogenous TCRα chain (TRAC) simultaneously. Freshly isolated PBMCs were stimulated with anti-CD3/CD28/CD2 for three days after which the cells were transiently transfected, in order to minimize off-target effects, with CRISPR ribonuleoprotein (RNP) complexes specific for PTPN22 and TRAC. This approach of transfecting protein complexes rather than DNA or RNA constructs avoids integration of foreign nucleic acid into the genome and ensures the targeting mixture is present for a short time only within the cells. Gene disruption frequencies were analyzed at the protein level in an untransfected cells, cells that were transfected with Cas9 alone (Cas9), cells that received Cas9 and the TRAC guide RNA (control) and cells that were targeted with multiplex genome editing of the *TRAC* and *PTPN22* locus (PTPN22^KO^).

CRISPR/Cas9 targeting achieved up to 90% reduction of PTPN22 protein expression analyzed by western blot (Fig. 1A). In agreement with the western blot, flow-IP analysis, in which protein from cell lysates was captured by antibody on beads and analyzed by flow cytometry, confirmed up to 90% reduction of PTPN22, while protein abundance of the internal control protein, AKT, remained unchanged (Fig. 1B). Additionally, gene disruption efficiency was verified at a genomic level for the *PTPN22* locus and two *in silico* predicted off-target regions (*chr.10, chr.16)* using the CHOPCHOP webtool [27] (Fig. 1C), which confirmed a specific gene disruption efficiency of up to 90% at the genomic level. Gene disruption efficiencies at the two off-target loci were usually below 10%, which is similar to the background detection of the assay as it was comparable to frequencies observed in non-targeted cells. *TRAC* disruption was analyzed at the protein level by FACS staining of cell surface TCRαβ chains. Up to 90% disruption of the TCRαβ expression was achieved in CD4 or CD8 T cells (Fig. 1D). In summary, we succeeded in efficient disruption of both the PTPN22 and the endogenous TCR expression using multiplex CRISPR-mediated genome editing.

**Figure 1.**
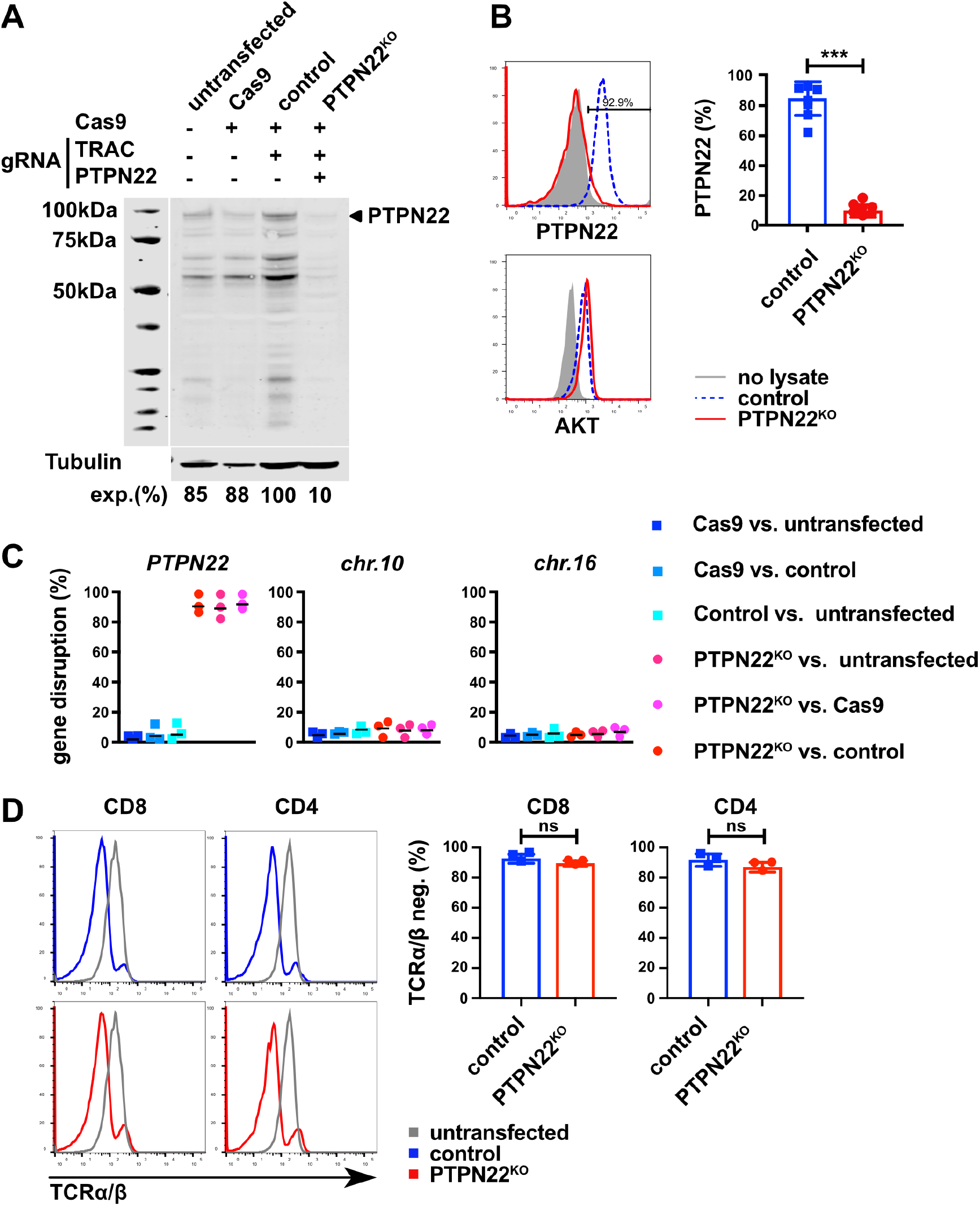
Multiplex genome editing targets PTPN22 and the TCR in human T cells. (A) PTPN22 knockout efficiency was analyzed by Western blot. Lysate equivalent to 2×10^6^ cells was run on the gel per well. The blot was probed with anti-PTPN22 and anti-tubulin antibodies. Data are shown from a representative experiment. (B) PTPN22 knockout efficiency was evaluated from lysates of T cells from 7 individual donors using Flow-IP. (C) Gene disruption frequencies for PCR products of the PTPN22 locus and two of the PTPN22 gRNA predicted off-target loci (chr.10 and chr.16) were determined using the CHOPCHOP webtool. Data are summarized from 3 healthy donors. (D) TCR knockout efficiency was evaluated by cell surface staining and flow cytometry in samples that received Cas9 and the TRAC guide RNA (control) or both TRAC and PTPN22 guide RNAs (PTPN22^KO^). Data are summarized from 3 healthy donors. Significance was determined using two-way ANOVA with Tukey’s post test for multiple comparisons. *\*P < 0.05, **P < 0.01, ***P < 0.001, ns = not significant*

### PTPN22 deletion does not alter the phenotype of EBV LMP2-specific T cells

T cells can be redirected to recognize TAAs by introduction of an exogenous TCR specific for the tumor antigen. PTPN22^KO^ and control T cells were transduced with a TCR specific for the HLA-A2-restricted EBV LMP2 peptide (CLGGLLTMV) that is frequently expressed in cancers including nasopharyngeal carcinoma, gastric cancer, Hodgkin lymphoma, and Burkitt lymphoma [28, 29]. For this TCR, the human TCRα and β variable fragments are linked to murine constant regions to enhance expression in human T cells and reduce mis-pairing with the endogenous TCR chains [30, 31]. After viral infection, cells were expanded in IL-7 and IL-15 because previous reports showed that these conditions favor the development of central memory T cells that were more efficient in killing tumors *in vivo* [32]. Seven days post transduction, cells were stained with Dextramers specific for the LMP2-TCR and an antibody against the mouse constant region. About 40% of the control and PTPN22^KO^ CD8 and CD4 T cells expressed the LMP2-TCR (Fig. 2A) and these TCR^+^ cells were enriched by magnetic bead separation for further functional analysis. Flow cytometry staining confirmed more than 90% purity of the enriched cells (Fig. 2B) which were re-stimulated with the antibody against the mouse constant region and anti-CD28, and expanded in IL-7 and IL-15.

**Figure 2.**
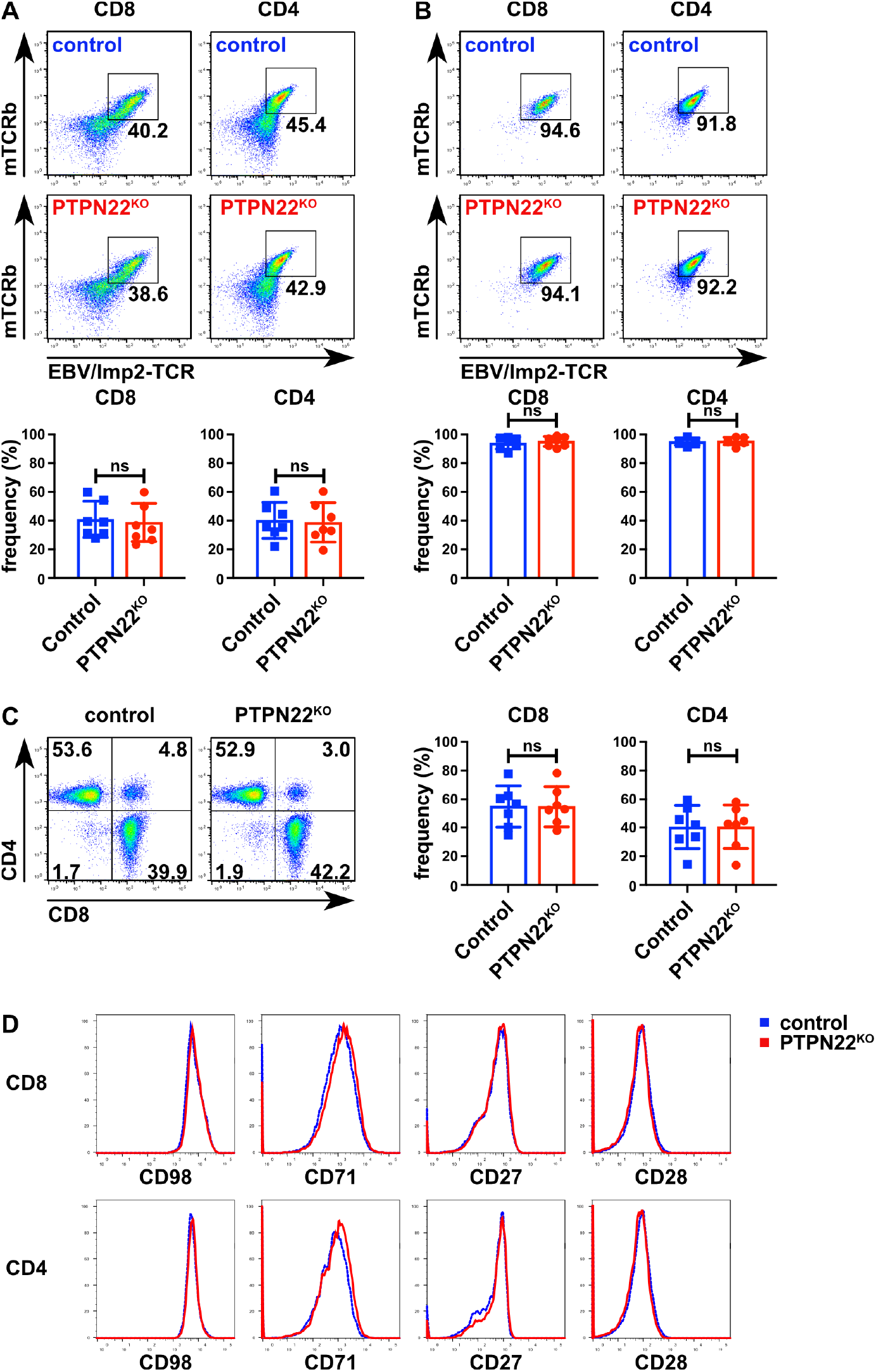
EBV/LMP2-specific PTPN22^KO^ and control T cells have similar CD8/CD4 frequencies and expression of surface markers. (A-B) TCR expression was determined before (A) and after (B) magnetic enrichment using antibodies against mouse TCRβ chain. Bar charts below show data summarized from 7 donors. (C) Representative dot plots and bar charts of data pooled from 7 donors show the frequencies of CD8 and CD4 T cells in control and PTPN22^KO^ samples. (D) Expression of CD98, CD71, CD27, and CD28 was evaluated by cell surface staining and flow cytometry. Data are shown from one representative experiment from 3 different donors (two other donors are shown in Figure S1). Significance was determined using two-way ANOVA with Tukey’s post test for multiple comparisons. *ns = not significant*

We next analyzed whether deletion of PTPN22 in human T cells changed the relative proportions of CD8 and CD4 T cells after three weeks of cell culture. We found that these proportions were unchanged, with both control and PTPN22^KO^ cells showing a slightly higher but comparable frequency of CD4 to CD8 T cells (Fig. 2C). Given that T cells often compete for nutrients in the tumor microenvironment and low nutrient concentrations can decrease T cell proliferation and cytokine secretion [33], we analyzed the expression of nutrient transporters by extracellular staining of the amino acid transporter CD98 and the transferrin receptor CD71. No difference was detected for CD98 and CD71 expression between PTPN22^KO^ and control cells (Fig. 2D and Suppl. Fig. 1). Analysis of the expression of the co-stimulatory molecules CD27 and CD28 also showed no difference (Fig. 2D and Suppl. Fig. 1). In summary, we were able to efficiently generate PTPN22^KO^ T cells that expressed the EBV LMP2-specific TCR. Deletion of PTPN22 did not alter the CD8/CD4 ratio or the expression of T cell surface markers after *in vitro* expansion.

### PTPN22^KO^ T cells are more responsive to tumor cells in vitro

Murine PTPN22^KO^ T cells were reported to have a proliferative advantage and were more cytotoxic than their wildtype counterparts [34, 35]. During *in vitro* expansion in IL-7 and IL-15, human T cells were counted by hemocytometer and viability was measured by trypan blue exclusion and hemocytometer counting. For all three donors shown, PTPN22^KO^ T cells proliferated more than the control cells (Fig. 3A) while viability was unaffected (Suppl. Fig. 2A). When the cells were tested in an *in vitro* cytotoxicity assay, human PTPN22^KO^ CD8 T cells were slightly more efficient in killing K562 tumor cells that expressed the LMP2 antigen presented on HLA-A2, in contrast to the control CD8 T cells from the same donor (Fig. 3B).

**Figure 3.**
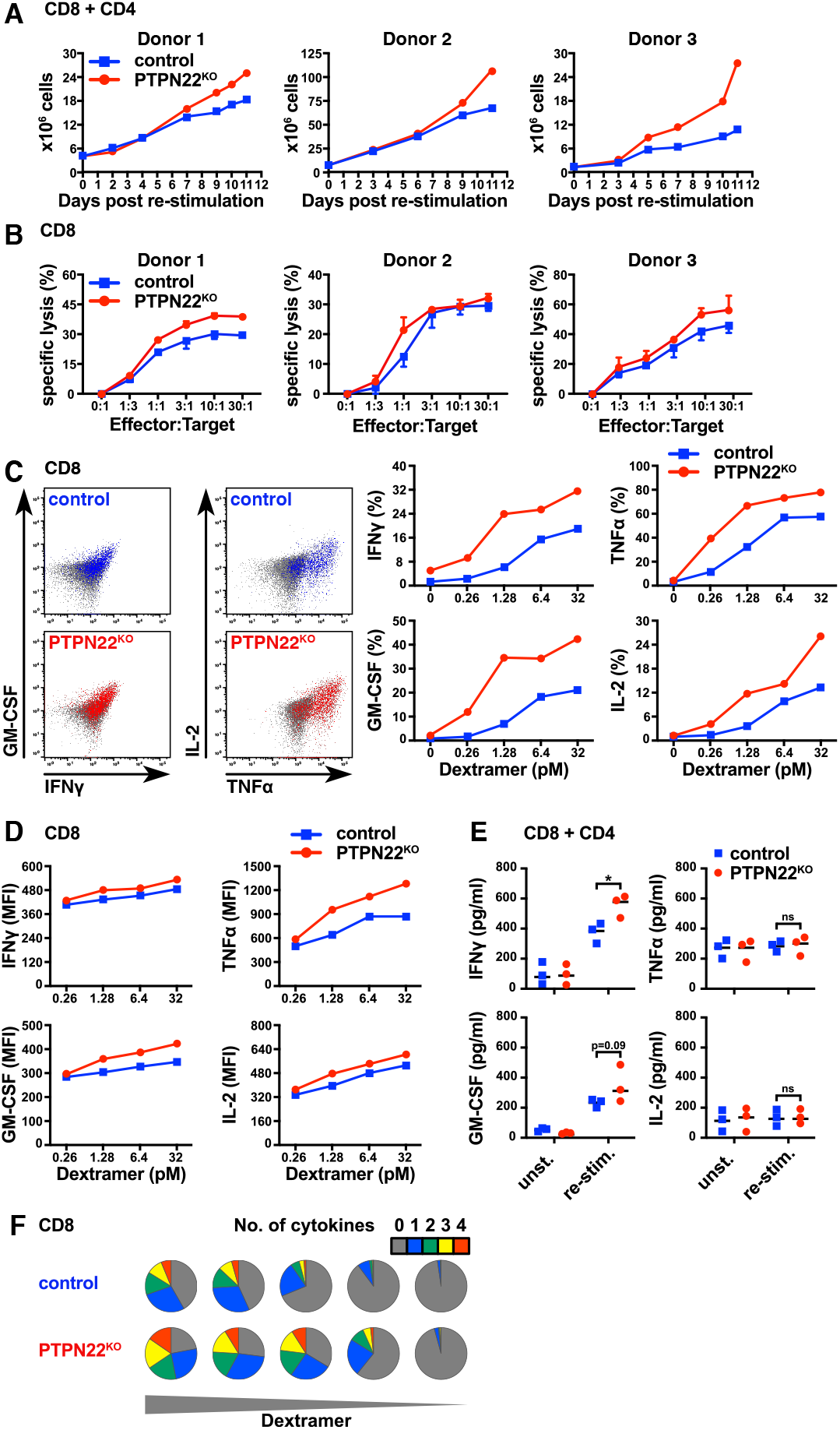
Human PTPN22^KO^ T cells proliferate more efficiently and are more functional. (A) Cell numbers were determined during in vitro expansion by hemacytometer counts. Data from 3 different donors are shown. (B) Specific lysis of K562 cells expressing the LMP2 antigen (Target) was analyzed after co-culture with increasing numbers of control or PTPN22^KO^ T cell (Effector) for 4 hours. Mean and SD of triplicate measurements are shown from 3 different donors. (C-D) PTPN22^KO^ and control T cells were re-stimulated for 4 hours with increasing concentrations of specific Dextramers. Frequencies of IFNγ^+^, TNFα ^+^, GM-CSF^+^ and IL-2^+^ T cells (C) and the mean fluorescence intensities (MFIs) of the cytokine-positive cells (D) were evaluated by intracellular staining and flow cytometry. Data are shown from one representative donor of 3 different healthy donors (two other donors are shown in Figure S2) (E) PTPN22^KO^ and control cells were co-cultured with media or with K562 cells expressing the LMP2 antigen. After 48 hours supernatants were collected and IFNγ, TNFα, GM-CSF and IL-2 secretion was analyzed by ELISA. Data of 3 different healthy donors are shown. Significance was determined using two-way ANOVA with Tukey’s post test for multiple comparisons. *\*P < 0.05, ns = not significant* (F) Polyfunctional cytokine expression for CD8 T cells was calculated using FlowJo software and visualized with Pestle and SPICE 6 software. Pie chart fragments represent the fraction of cells expressing the number of cytokines indicated in the key. Data are shown from one representative donor of 3 different healthy donors (two other donors are shown in Figure S2).

Tumors have various strategies to evade immune cell recognition including downregulation of the expression of peptide:MHC antigens [13]. To recapitulate low antigen concentrations in the tumor microenvironment, T cells were re-stimulated with decreasing concentrations of LMP2-specific dextramers in the presence of constant concentrations of anti-CD28 and cytokine expression was measured by intra-cellular (IC) staining and flow cytometry. PTPN22^KO^ CD8 T cells retained higher frequencies of IFNγ^+^, TNFα ^+^, GM-CSF^+^ and IL-2^+^ cells upon re-stimulation throughout the varying dextramer concentrations (Fig. 3C and Suppl. Fig. 2B). Furthermore, PTPN22^KO^ CD8 T cells expressed more of all measured cytokines on a per cell basis as assessed by mean fluorescence intensity (MFI) (Fig. 3D and Suppl. Fig. 2C).

In addition to stimulation with pMHC dextramers we also assessed cytokine production in PTPN22^KO^ and control T cells following 48h co-culture with K562 tumor cells that expressed the TCR specific tumor antigen, LMP2, plus HLA-A2. As seen with dextramer restimulation, the PTPN22^KO^ cells secreted more IFNγ and GM-CSF as measured by ELISA in response to tumors (Fig. 3E). However, in contrast to the intracellular cytokine staining, no TNFα and IL-2 secretion was detected in response to LMP2-expressing tumor cells from either control or PTPN22^KO^ T cells. This apparent discrepancy between the IC staining and ELISA data could be accounted for by the different duration of the re-stimulation between these experiments, namely 4h for the intracellular staining versus 48h for the ELISA assay. During the 48h culture with tumor cells, the cells themselves may have consumed the IL-2 and TNFα cytokines for which they express receptors. In summary, we showed that human PTPN22^KO^ T cells have a proliferative advantage, express higher amounts of inflammatory cytokines and are more efficient in killing tumor cells *in vitro* than cells from the same donor which express PTPN22.

T cells that express multiple cytokines simultaneously are called polyfunctional and it was shown that polyfunctional T cells are more effective in tumor elimination [16, 17]. Thus, we compared frequencies of polyfunctional cells between the control and PTPN22^KO^ CD8 T cells. To demonstrate polyfunctionality, frequencies of cells, as measured by FACS analysis, that expressed 0, 1, 2, 3, and 4 cytokines simultaneously were calculated for the different dextramer concentrations. Already at low antigen concentrations the PTPN22^KO^ CD8 T cell population contained higher frequencies of cells that expressed more than two cytokines compared to the control cells (Fig. 3F and Suppl. Fig. 2D). This suggests that PTPN22^KO^ CD8 T cells have the ability to retain polyfunctionality better in environments with low antigen concentrations, such as might be anticipated within the suppressive tumor microenvironment.

### PTPN22^KO^ T cells preserve more polyfunctional responses when antigen is limiting

In order to test whether the improved responses gained from knocking out PTPN22 applied to other TCR specificities with different affinities, we transduced control and PTPN22^KO^ T cells with a second TCR specific for the HTLV Tax peptide (LLFGYPVYV). The HTLV Tax-specific TCR was reported to have approximately a 10-fold higher affinity and longer half-life (t_1/2_) for pMHC interaction (K_D_=2μM, t_1/2_=6.7sec) compared to the EBV LMP2 TCR (K_D_=23μM, t_1/2_=2.3sec) [36]. When PTPN22^KO^ and control T cells were re-stimulated with peptide:MHC dextramers specific for the Tax-TCR, they also secreted more IFNγ and GM-CSF as measured by ELISA than the PTPN22 expressing control cells (Fig. 4A) These data demonstrate that deletion of PTPN22 improved cytokine production for at least two, EBV LMP2 and HTLV Tax, distinct TCR-transduced primary T cells which differ in affinity by 10-fold.

**Figure 4.**
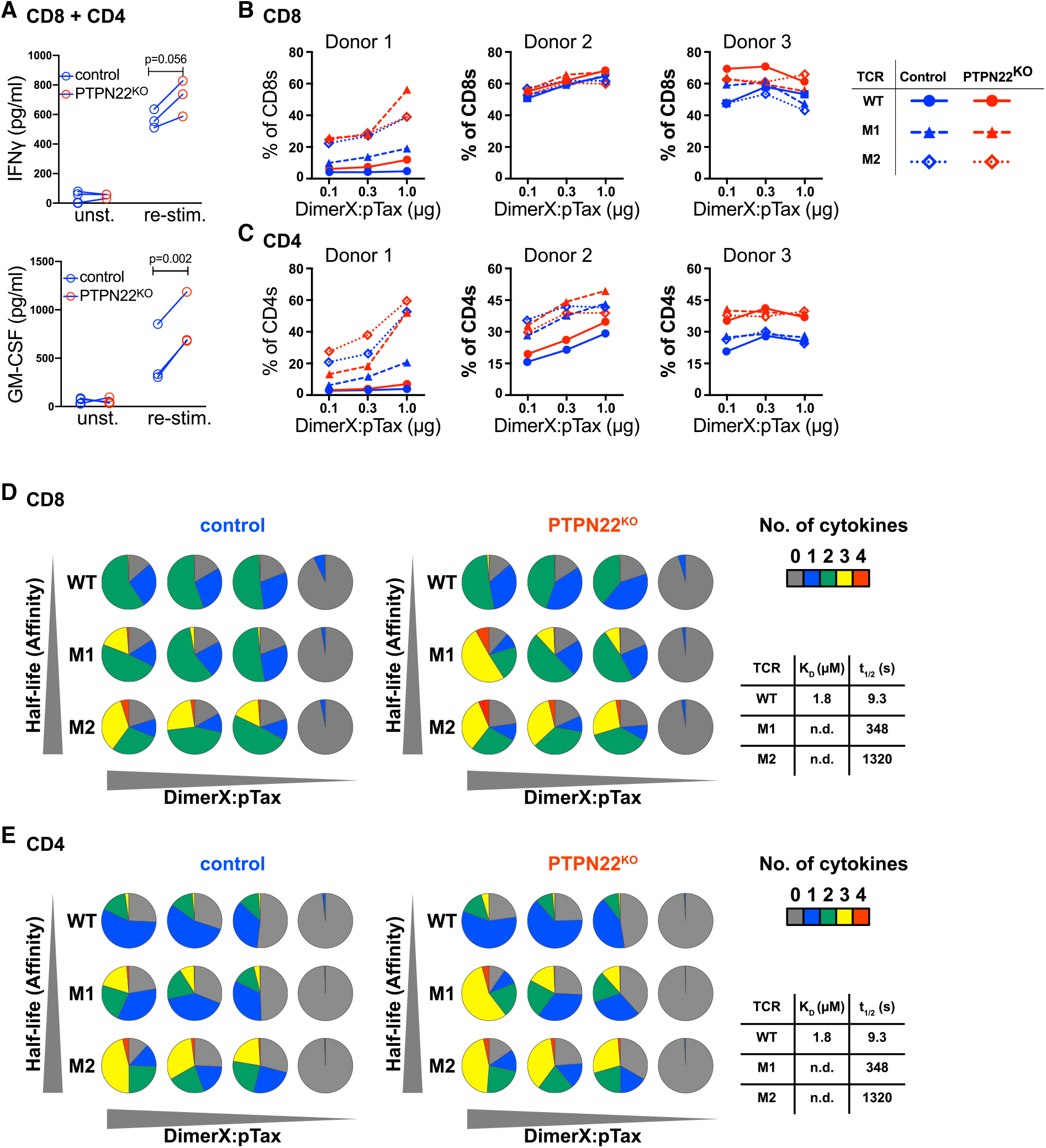
PTPN22KO T cells remain more polyfunctional after re-stimulation. (A) PTPN22KO and control cells transduced with Tax-TCR and were re-stimulated with 25pM of pTax:HLA-A2 Dextramers. After 48 hours supernatants were collected and IFNγ GM-CSF and secretion was analyzed by ELISA. Data of 3 different healthy donors are shown. Significance was determined using a paired t-test. (B-E) PTPN22KO and control T cells were transduced with Tax-TCRs that have increasing affinities for the Tax antigen (WT<M1<M3) and re-stimulated with increasing concentrations of DimerX+pTax for 4 hours. Frequencies of GM-CSF+ cells for CD8 (B) and CD4 (C) T cells were evaluated by intracellular staining and flow cytometry for 3 different donors. Polyfunctional cytokine expression for CD8 (D) and CD4 (E) T cells from donor 1 was calculated using FlowJo software and visualized with Pestle and SPICE 6 software. Pie chart fragments represent the fraction of cells expressing the number of cytokines indicated in the key. Data are shown from one representative donor of 3 different healthy donors (two other donors are shown in Figure S3).

A concern with adoptive T cell therapy is that, in general, tumor Ag-specific TCRs are of low affinity and thus stimulate poor effector responses. However, increasing the affinities of TCRs for tumor antigens beyond the natural affinity range, not only risks breaking tolerance, but was also shown paradoxically to result in loss of sensitivity to peptide [37]. To ask how removing PTPN22 influenced the response of T cells with different affinity peptides in the particular context of improving polyfunctionality of cytokine production and therefore anti-tumor efficacy, we obtained several constructs for the Tax-TCR which had been engineered and selected by phage display to have altered binding properties for the peptide:MHC [38]. We used three TCRs specific for Tax: HLA-A2 that were shown to have a range of peptide:MHC-binding half-lives which were measured as: WT t_1/2_ 9.3sec, M1 t_1/2_ 348sec and M2 t_1/2_ 1320sec. The higher affinity TCRs had been shown to confer faster responsiveness to peptide in primary transduced T cells (M2>M1>WT), but also decreased sensitivity to lower concentrations of antigen (WT>M1>M2) [39]. Primary PBMC were stimulated as before and CRISPR/Cas9 used to delete TRAC ± PTPN22 before transduction with the specific TCRs. In order to limit the multivalency of the antigen we re-stimulated the T cells with increasing concentrations of DimerX-pTax complexes (0.1μg, 0.3μg, 1μg) to mimic different antigen availability and measured cytokine production.

Four functional outputs, IFNγ, TNFα, GM-CSF, and IL-2, were measured in response to cognate antigen exposure by intracellular staining of both CD4 and CD8 TCR expressing cells. Detailed analysis of GM-CSF production by CD8 (Fig 4B) and CD4 (Fig 4C) T cells from three donors transduced with the WT, M1 and M2 TCRs in response to a titration of DimerX-pTax complexes, are shown. Overall loss of PTPN22 increased the response of cells expressing each of the TCRs until the maximum response was reached, at which point no further improvement in cytokine production was observed (e.g. Fig 4B, donor 2). For both CD4 and CD8 T cells from these donors the higher affinity receptors progressively induced greater proportions of cytokine producing cells, M2>M1>WT, and this held true in both PTPN22 sufficient and deficient T cells.

However, when we assessed the polyfunctionality of the cells we found that in PTPN22 expressing control CD8 and CD4 T cells, there was a clear correlation between the affinity of the TCR and the propensity to produce multiple cytokines. Thus, higher affinity TCRs induced progressively greater numbers of polyfunctional cells at any given peptide concentration, whereas the cells expressing the lower affinity WT TCR were more limited in their range of cytokine production at all concentrations of pMHC (Fig. 4D and 4E and Suppl. Fig. 3). This was different for the PTPN22^KO^ CD8 and CD4 T cells. PTPN22^KO^ T cells transduced with the intermediate affinity M1-TCR showed greater polyfunctionality at all pMHC concentration compared to both M1 TCR transduced PTPN22-sufficient control cells and to PTPN22^KO^ cells expressing either lower (WT) or higher (M2) affinity TCRs. For the lower and higher affinity TCRs loss of PTPN22 had little impact on polyfunctionality compared to PTPN22-sufficient cells with only a marginal increase the range of cytokines produced. In conclusion, PTPN22^KO^ T cells transduced with a low affinity TCR specific for the Tax antigen secreted more IFNγ and GM-CSF. Frequencies of GM-CSF^+^ and IL-2^+^ T cells were increased after PTPN22 deletion. Additionally, increasing TCR affinity when PTPN22 was also deleted, resulted in improved polyfunctionality in response to lower antigen doses, indicating that removal of PTPN22 would be beneficial over a number of different affinity TCRs.

### PTPN22^KO^ T cells are more efficient in controlling tumor growth in vivo

Human PTPN22^KO^ T cells were more cytotoxic and more polyfunctional in response to antigen stimulation *in vitro*. This is similar to what we found with mouse PTPN22^KO^ T cells which, furthermore, had enhanced anti-tumor activity *in vivo* [26]. To test whether human PTPN22^KO^ T cells also were more efficient in eliminating tumor cells *in vivo*, we injected NSG mice with K562 tumor cells that expressed HLA-A2 plus LMP2 together with luciferase (Fig. 5A). Once tumors became palpable, control or PTPN22^KO^ T cells that express the EBV LMP2-specific TCR were injected intravenously.

**Figure 5.**
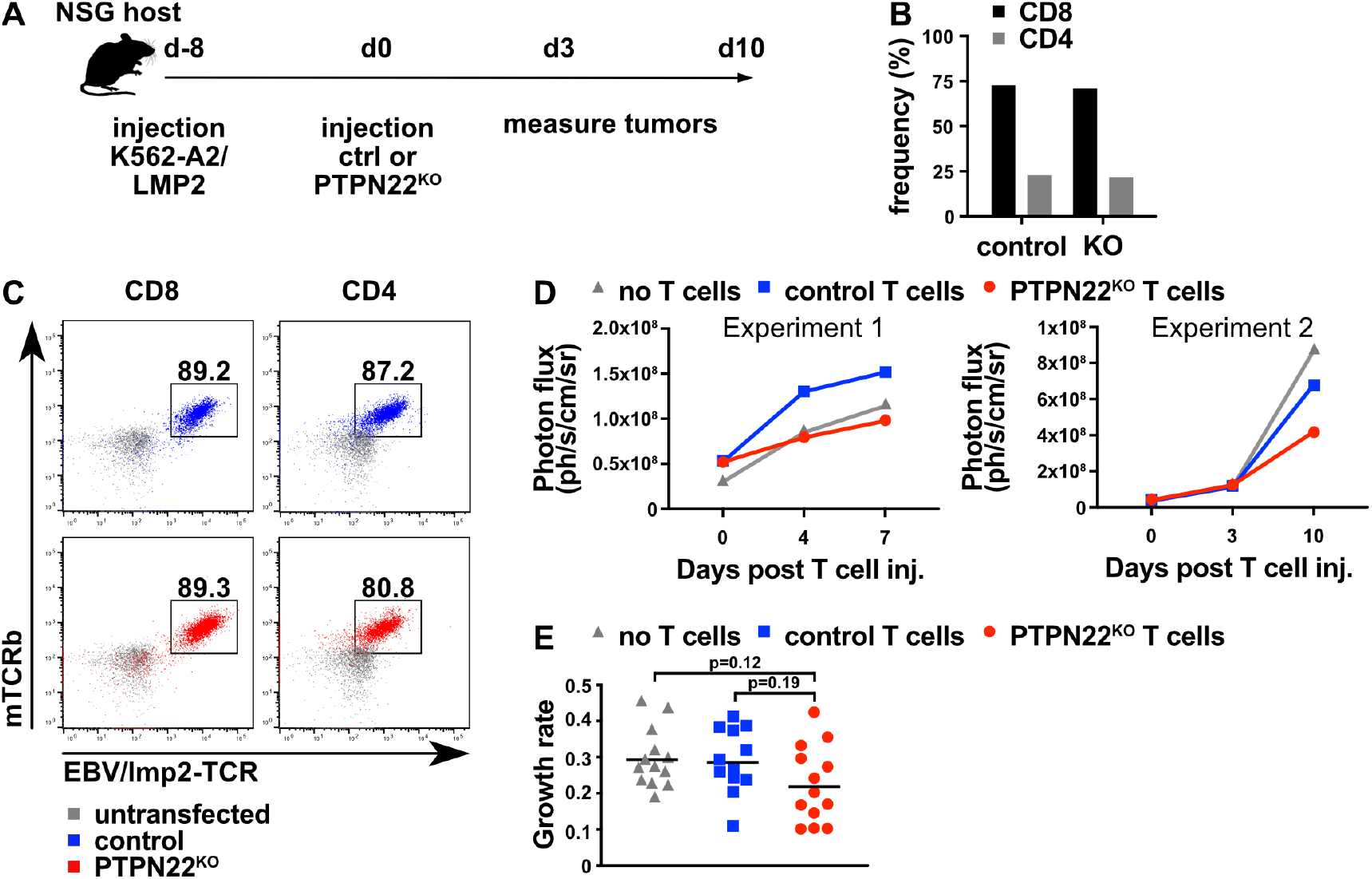
PTPN22^KO^ T cells are more efficient in controlling tumor growth. NSG mice were injected s.c. with 2×10^6^ K562 cells expressing HLA-A2 and the LMP2 antigen. After 8 days, 10^6^ EBV/LMP2-TCR transduced PTPN22^KO^ or control cells were injected i.v. (A) Scheme of experimental design. (B) CD8 and CD4 T cells were mixed at the same ratio (4:1) before adoptive transfer and analyzed for CD8 and CD4 expression using flow cytometry. (C) PTPN22^KO^ and control T cells were evaluated for EBV/LMP2-TCR expression using flow cytometry before adoptive transfer. (D) Mice were assessed for tumor establishment on day 0 using bioluminescence imaging. All mice were assessed for tumor growth on day 4 and 7 (Experiment 1) or day 3 and 10 (Experiment 2) after T cell injection. Graphs show tumor growth over time using photon flux. Data are shown from two representative experiments using 5-8 mice per group. (E) Graph shows the growth rate of both experiments. Significance was determined using two-way ANOVA with Tukey’s post test for multiple comparisons.

The importance of CD4 T cell help for benefitting adoptive T cell therapy has been shown in multiple studies [30, 40, 41]. Thus, we decided to inject mice with a mixture of CD8 and CD4 T cells. In order to ensure equivalent CD8 and CD4 frequencies amongst both the control and PTPN22^KO^ T cells, we first separated the CD8 and CD4 T cells by MACS depletion and then remixed the cells at a precise 4:1 ratio (Fig. 5B). Both PTPN22^KO^ and control T cells expressed similar frequencies and intensities of the EBV LMP2-specific TCR. Flow cytometry revealed that more than 80% of the injected T cells were EBV/LMP2-TCR positive (Fig. 5C). Tumor growth was monitored using bioluminescence over several days after mice received the T cells and the photon flux over time for two independent experiments are shown (Fig. 5D). Calculating the growth rate and amalgamating the data from both experiments showed a clear trend for the PTPN22^KO^ cells to better control tumor growth in vivo, although the data did not reach significance given the large variance among the individual mice (Fig. 5E). Overall these data indicate that human PTPN22^KO^ T cells are more efficient than PTPN22-sufficient T cells from the same donor in controlling tumor growth *in vivo,* and parallel our findings with mouse PTPN22^KO^ T cells [26].

## Discussion

Immunotherapies aimed at harnessing T cells to destroy tumor cells have demonstrated remarkable results in some cancer patients [1–5], but have not been so successful in others. A major issue remains the ability of tumors to suppress immune responses, therefore, gaining a greater understanding of mechanisms that negatively influence T cell function is critical for improving the role of T cells in cancer immunotherapy. In this study we describe the potential value of disrupting a negative regulator of TCR signaling, PTPN22, to improve the efficacy of human adoptive T cell therapy, complementing our published findings with mouse PTPN22^KO^ T cells [25, 26, 42]. CRISPR-mediated deletion of PTPN22 in human T cells resulted in higher secretion of the inflammatory cytokines IFNγ and GM-CSF. Additionally, after re-stimulation of human PTPN22^KO^ T cells we detected greater numbers of polyfunctional cells, expressing two or more cytokines, which is thought to be beneficial for tumor control. PTPN22^KO^ T cells were also more cytotoxic *in vitro* and superior in controlling tumor growth *in vivo* in a murine transfer model.

PTPN22 can dephosphorylate LCK, ZAP70, TCR*ζ* and VAV1 thereby limiting TCR proximal signaling [43]. Therefore, removal of PTPN22 can increase the sensitivity of the T cell response to antigen which may be beneficial when trying to optimize responses to weak tumor-specific antigens. TCR signaling is dependent on the affinity of the TCR and is defined by a combination of the strength of binding to a pMHC molecule combined with the on and off rates. However, a more useful parameter for whether an TCR:pMHC interaction will stimulate a T cell response is the functional avidity which measures responsiveness as an outcome and is influenced by a number of factors including TCR affinity, stability, expression levels of the TCR and the coreceptors, distribution and composition of the signaling molecules, and expression levels of molecules that attenuate T cell function [44–47]. Functional avidity impacts upon T cell functions such as cytokine release, cytotoxicity, and antitumor responses [46]. There is substantial evidence that better T cell function is associated with higher TCR affinity *in vitro* and *in vivo* [47–49], and higher TCR affinities also improve functional avidity, but only until a certain threshold [39, 47]. Murine PTPN22 was shown to play a particular role in limiting the response of OT-1 TCR transgenic T cells to weak peptide ligands, which was enhanced in PTPN22^KO^ T cells, but had less influence on the response to strong antigens [34]. Thus, in murine T cells PTPN22 appears to contribute to the functional avidity of the T cell, most likely by influencing TCR proximal signaling molecules. In this study we addressed how PTPN22 deletion impacted on the functional avidity of human T cells by examining responses elicited by TCRs of various affinities to a common ligand, the Tax peptide from HTLV. A surprising finding was that a substantial impact of PTPN22 deletion was not only that individual cytokine production was amplified, but also that the number of individual cytokines produced per cell was expanded. Thus, removal of PTPN22 seemed to lower the TCR affinity needed to trigger a polyfunctional response, except when cells expressed the highest affinity Tax-TCR (M2) whereupon fewer T cells expressed more than two cytokines simultaneously. Although PTPN22 deficiency decreased the affinity needed to elicit a polyfunctional response, an optimal window of TCR affinity still existed. In this context, deleting PTPN22 in T cells expressing CARs might not be beneficial given the high affinity binding of CARs compared to TCRs and the tendency of CAR expressing T cells to veer towards excessive cytokine production potentially causing cytokine release syndrome.

We used a mixture of CD8 and CD4 T cells in the *in vivo* tumor model. Successful re-direction of CD4 T cell effector activities against MHC class I (MHCI)-restricted epitopes has been shown previously [30, 50]. Since the transfected TCRs were MHCI-restricted, we expected that the lack of the CD8 co-receptor might impair the functional avidity of the engineered CD4^+^ T cells, given that the absence of CD8 would increase the TCR affinity needed to reach the same functional avidity [30]. Our experiments confirmed that at lower TCR affinities (WT and M1 Tax-TCRs), CD4 T cells either express no cytokines or only one cytokine, whereas CD8 T cells already show a reasonable percentage of dual cytokine producers (Fig. 4D and E). At higher affinities (M2) the polyfunctionality between CD4s and CD8s was comparable. PTPN22 deficiency in CD4 T cells achieved simultaneous expression of several cytokines at lower TCR affinities and lower antigen concentrations compared to control CD4 T cells. This suggests that better functionality of PTPN22^KO^ cells is not limited to CD8 T cells but can also improve the function of antigen-redirected CD4 T cells. Since human PTPN22^KO^ T cells retained a more polyfunctional cytokine expression upon TCR re-stimulation (Figure 3F and 4D-E), they might be able to remain more functional when exposed to the suppressive tumor environment and be rendered less exhausted than PTPN22 sufficient T cells. Further study is warranted to specifically address this hypothesis.

In addition to their polyfunctionality, one mechanism for superior control of tumor growth by human PTPN22^KO^ T cells may be linked to their production of greater quantities of inflammatory cytokines including GM-CSF, TNFα and IFNγ . However, cytokines can have diverse effects with mixed outcomes on cells in the tumor microenvironment and therefore tumor growth. GM-CSF can induce anti-tumor immunity by mainly acting on antigen presenting cells [51, 52]. Additionally GM-CSF can induce tumor progression and stimulate the expression of PD-L1 on tumor cells which in turn may increase the proliferation and activation of myeloid-derived suppressor cells in the tumor [53–56]. IFNγ has pleiotropic functions and was shown to induce apoptosis and to increase antigen presentation on tumor cells [11, 57, 58]. Controversially, IFNγ can promote tumor growth by upregulating inhibitory molecules [59]. TNFα was shown to induce tumor and endothelial cell death [60–62]. Additionally, TNFα-dependent processes were associated with increased expression of the immune checkpoint molecule CD73 and loss of antigen expression in melanoma leading to poor recognition by antigen-specific T cells [63, 64]. It is possible that the concentrations of intra-tumoral cytokines in particular are important for the outcome of the cancer therapy. For instance, it was recently shown that CD8^+^ T cells that used TNFα as part of their anti-tumor function slowed the progression of MC38 colon carcinomas in perforin-deficient mice in the context of anti-PD-1 therapy [65]. In another study, adoptively transferred CD4 T cells into mice with established colorectal cancers [66] shaped tumor metabolism through TNFα which synergized with chemotherapy to induce tumor cell oxidative stress. In both studies, TNFα production by T cells was not enough to reject the tumors and only upon treatment with anti-PD-1 or after chemotherapy was the effect of TNFα apparent. It might be that in general physiological intra-tumoral TNFα levels are insufficient to induce cancer regression [67]. Therefore, increased expression by PTPN22^KO^ T cells might produce sufficient TNFα abundance in the tumor to induce toxicity. Overall, although there was a clear trend for transferred human PTPN22^KO^ T cells to better restrain tumor growth *in vivo* compared to control T cells from the same donor, the data did not reach significance given the high variance in tumor growth in individual mice. The pleiotropic nature of cytokines and the varying effects at different cytokine doses may in part explain this variability. Additional experiments are necessary to clarify the role of cytokine-mediated killing by PTPN22^KO^ cells and to assess whether combining strategies such as PTPN22 deletion with checkpoint inhibitors would be able to induce a more potent tumor elimination.

Other studies have applied the strategy of targeting of intracellular phosphatases in T cells and these have shown some promising results to improve adoptive T cell therapy of tumors. Examples include the dual-specific phosphatase PAC1/DUSP2, which was found to negatively regulate STAT3 signaling and is involved in Th17 differentiation [68], and when knocked out in T cells resulted in higher expression of inflammatory molecules and superior elimination of tumor cells [69]. DUSP2 seemed to restrain effector function in T cells, as shown in a LCMV infection model, where DUSP2 deficient T cells expressed fewer inhibitory molecules and were more functional suggesting that deletion of DUSP2 might result in less exhausted T cells in the tumor microenvironment. Similarly, PTPN2 deletion in Her2-specific CAR T cells was shown to increase efficiency of tumor eradication [70], while another study showed that CRISPR-mediated PTPN2 deletion in T cells attenuated tumor growth [71]. PTPN2 negatively regulates TCR signaling by dephosphorylating the TCR proximal kinase LCK [72] and its deletion lowers the threshold for T cell proliferation and enhances CD8 T cell responses to low affinity antigens. Additionally, PTPN2 is a negative regulator of JAK/STAT signaling pathways downstream of cytokines [73]. Finally, adoptively transferred PTPN6/SHP-1 deficient T cells demonstrated enhanced effector function and tumor clearance [74]. SHP-1, was suggested to negatively regulate TCR signaling by dephosphorylating LCK and thus set the threshold of activation [75–77]. Moreover, SHP-1 was proposed to interact with the inhibitory receptor, PD-1, thereby negatively influencing TCR signaling [78, 79]. Thus, as a more general strategy, deletion of phosphatases might be used to lower the TCR activation threshold to respond to low affinity antigens while additionally targeting cytokine signaling pathways thereby enhancing T cell function. Overall these studies demonstrate that targeting phosphatases has interesting potential for improving T cell therapy and several other phosphatases have been suggested as possible targets [80].

In conclusion, we showed that deletion of PTPN22 in human T cells increased their cytokine expression and cytotoxicity and made the cells more effective in controlling tumor growth. PTPN22^KO^ T cells seemed to be more polyfunctional at low antigen concentrations. This suggests that they might be more functional within the suppressive tumor microenvironment thereby overcoming the limitations of immunotherapy for solid tumors.

## Materials and Methods

### Human PBMCs

Peripheral blood samples were collected at the University of Edinburgh. Collection of blood samples was approved by the institutional ethics committee and samples were collected following informed written consent. Lithium heparin containing vacutainers (green top, BD Biosciences, USA) were used for plasma collection, immediately transferred into Leucosep tubes (Greiner Bio-one, Germany), centrifuged at 1000g for 15 minutes, and PBMCs collected. Samples were either used directly for experiments or frozen at –80°C until needed. PBMCs were activated for 3 days with anti-CD3/28/2 reagent following manufacturer’s instructions (Stemcell, Canada) in complete medium consisting of Iscove’s Modified Dulbecco’s Medium (IMDM, Thermo Fisher Scientific, USA) supplemented with 10% FCS, L-glutamine and antibiotics and containing 10ng/mL recombinant human IL-7 and IL-15 (Peprotech, USA).

### Animals

NOD.Cg-Prkdc^scid^ Il2rg^tm1Wjl^/SzJ (NSG) mice were bred and maintained in pathogen-free conditions at the University of Edinburgh animal facilities in accordance with the UK Home Office and local ethically approved guidelines, and used as recipients of human tissue and cell transplantation at 6 to 9 weeks of age.

### Cells and cell lines

K562-A2-luc2 and K562-A2/LMP2-GFP-luc2 cells were used for *in vitro* and *in vivo* experiments. Phoenix-Ampho cell lines were used for retroviral transductions. K562 and Phoenix-Ampho cell lines were obtained from Hans Stauss (University College London). K562-A2/LMP2-GFP-luc2 cells stably express HLA-A2 and the EBV latent membrane protein 2 (LMP2) as a tumor-specific antigen, while K562-A2-luc2 only express HLA-A2. Phoenix-Ampho cells were used for retroviral transductions. All cell lines were cultured in complete medium and split every 2-3 days and routinely tested to ensure they were free of mycoplasma.

### CRISPR

Guide RNA sequences for PTPN22 (TTATCAGGGATAGTTCTACC) and TRAC guide RNA (CAGGGTTCTGGATATCTGT) were used for multiplex-CRISPR/Cas9. Cas9 ribonucleoprotein (RNP) complex were generated by incubating 2μL of 100μM tracrRNA and 2μL of 100μM crRNA in 25μL Nuclease-free buffer (Integrated DNA Technologies, USA) at 95°C for 5 min and then mixed with 2μL of 5mg/mL Truecut Cas9 protein (Thermo Fisher Scientific, USA) for 10 min at 37°C in a total volume of 50μL. RNP complexes were mixed directly prior to transfection and combined with 10^6^ activated PBMCs. Transfections were carried out using the Neon transfection system (Thermo Fisher Scientific, USA) following manufacturer’s instructions. The conditions used were: 1600V, 10ms, 3 pulses.

### Retroviral transduction

EBV/LMP2- and HTLV/Tax-specific TCRs have been described previously [30, 39]. After CRISPR-mediated knockout, T cells were rested for 1 hour in complete medium supplemented with 10ng/ml IL-7/IL-15 at 37°C. Then, 10^6^ cells per well were transferred into RetroNectin-coated 24well plates (Takara Bio, USA). Virus supernatant was centrifuged (300xg, 5min) and 500μl was transferred onto the PBMCs. The plate was centrifuged for 90min at 500xg at 32°C. Afterwards, the supernatant was replaced by complete medium containing 10ng/ml IL-7 and IL-15.

### Magnetic enrichment

Transduced T cells were labelled with anti-mouse TCRβ-Biotin (clone H57-597, Thermo Fisher Scientific, USA) for EBV/LMP2-TCR or anti-TCRVβ13.1-PE (clone H131, BioLegend, USA) for Tax-TCR for 15 min at 4°C. Afterwards, cells were enriched using the EasySep Release Human Biotin Positive Selection kit (Stemcell, Canada) for EBV/LMP2 according to manufacturer’s instructions. EasySep Release Human PE Positive Selection kit II (Stemcell, Canada) was used to enrich Tax-TCR+ cells according to manufacturer’s instructions.

### Western Blot

Cells were lysed in 1% TritonX-100, 0.5% n-dodecyl-b-D-maltoside, 50mM Tris-HCl, 150mM NaCl, 20mM EDTA, 1mM NaF, and 1mM sodium orthovanadate containing 1x Protease Inhibitor cocktail (Merck Group, Germany). Lysate equivalent to 2×10^6^ cells per well was run on a NuPAGE 4 to 12% Bis-Tris gel (Thermo Fisher Scientific, USA). Western blots were performed on samples and membranes were probed with anti-PTPN22 (clone D6D1H, Cell Signaling Technology, USA) and anti-tubulin (clone TU-02, Santa Cruz Biotechnology, USA). For detection, goat anti-mouse-A680 and goat anti-rabbit-IR800 (Thermo Fisher Scientific, USA) antibodies were used and visualization was performed using infrared imaging system (Odyssey; LI-COR Biosciences, USA).

### Flow-IP

5μM and 10μM CML latex beads (Thermo Fisher Scientific, USA) were coated respectively with anti-AKT (clone SKB1, Merck Group, Germany) or anti-PTPN22 (clone D6D1H, Cell Signaling, USA) antibodies in MES coupling buffer (50mM MES, 1mM EDTA, pH=6.0). Cells in exponential growth were lysed in IP lysis buffer (0.15M NaCl, 50mM TrisHCl, 1% Triton X-100, 0.5% n-Dodecyl-β-D-Maltoside, 1x Protease Inhibitor cocktail (Merck Group, Germany), 1mM Na3VO4, 1mM NaF) at a concentration of 100,000 cells/μL and incubated for 20min on ice followed by centrifugation at 10,000xg for 10min to remove nuclei. 6μL of the lysate was mixed with 9,000 or 12,000 beads and incubated for 2 hours at 4°C at 200rpm. Then, beads were washed with PBS and stained with anti-PTPN22 (anti-Lyp, R&D Systems, USA) for 20min at 4°C. This was followed by incubation with anti-goat IgG-FITC (Merck Group, Germany) and anti-AKT-AlexaFluor647 (clone 5G3, Cell Signaling, USA) for 20min at 4°C. Bead fluorescence was measured on a MACS Quant Analyzer 10 (Miltenyi Biotec, Germany).

### TIDE analysis

Genomic DNA was extracted from 2×10^5^ T cells using 20μl DNA lysis buffer (30mM Tris-HCl, 10mM EDTA, 0.1% SDS, 0.5% Tween 20). Cell suspensions were incubated at room temperature for 15 min, followed by two 5 min incubation steps at 55°C and 98°C. 1μl was used for the Phusion High-Fidelity DNA Polymerase PCR reaction (New England Biolabs, USA) following manufacturer’s instructions. PCR conditions used were: 95°C 3 min, 35 cycles of 95°C 1 min, 60°C 30 sec, 72°C 1 min, 72°C 10 min. Primers sequences used for the *PTPN22* gene (TGTTCTCACCTAGTCCTCCG & GTGGCTGAGAAGCCCAAGAAT) and for the off-target on chromosome 10 (TCCTCAATCAGGGCAAGGAAT & GGCATTGAGGACACTTCTTGTT) and chromosome 16 (GCCTCGTAGGCAAAGAAAAAGG & TATAGCTATCCCGGGGCTGT). The PCR products were purified from a 1% agarose gel using the Monarch DNA Gel Extraction kit (New England Biolabs, USA) and send for Sanger sequencing provided by Edinburgh Genomics. Sequences were analyzed using an online webtool (https://tide.deskgen.com).

### Re-stimulation experiments

EBV/LMP2-TCR transduced cells were re-stimulated with a titration of peptide-specific MHC Dextramer (0.8, 4, 20, 100pM, Immudex, Denmark) in the presence of 1μg/ml anti-CD28 (clone CD28.2, BioLegend, USA) and 10mg/ml Brefeldin A (Cayman Chemical, USA) for 4 hours at 37°C followed by intracellular staining of cytokines and analysis using flow cytometry. For cytokine secretion 0.2×10^6^ EBV/LMP2-TCR transduced cells were co-cultured with 0.02×10^6^ K562-A2/LMP2-GFP-luc2 cells for 48h and supernatants were collected. IFNγ, TNFα, GM-CSF, and IL-2 concentrations were analyzed using the respective LEGEND MAX Human ELISA Kits (BioLegend, USA).

DimerX (BD Biosciences, USA) was mixed with 160 molar excess of Tax peptide (Thermo Fisher Scientific, USA) and incubated for 24h at 37°C. Flat-bottom 96 well plated were coated with a titration of DimerX:pTax complexes in 50μl PBS for 24h at 4°C. Tax-TCR transduced cells were re-stimulated with DimerX:pTax complexes (pM) in the presence of 1μg/ml anti-CD28 and 10mg/ml Brefeldin A for 4 hours at 37°C followed by intracellular staining of cytokines and analysis using flow cytometry. For cytokine secretion 0.2×10^6^ Tax-TCR transduced cells were re-stimulated with 8pM MHC Dextramer (Immudex, Denmark) in the presence of 1μg/ml anti-CD28 for 48h and supernatants were collected. IFNγ, TNFα, GM-CSF, and IL-2 concentrations were analyzed using the respective LEGEND MAX Human ELISA Kits.

### Flow cytometry

10^5^ cells were stained for flow cytometry. Anti-CD8-PacificBlue (clone SK1, BioLegend, USA), anti-CD4-APC/eFluor780 (clone RPA-T4, Thermo Fisher Scientific, USA), anti-CD3-PerCP/Cy5.5 (clone OKT-3, Thermo Fisher Scientific, USA), anti-mouse TCRβ PacificBlue (clone H57-597, Thermo Fisher Scientific, USA), anti-TCRαβ-FITC (clone REA652, Miltenyi Biotec, Germany), anti-TCRVβ13.1-PE (clone H131, BioLegend, USA) were used for extracellular staining. Fixation buffer and Permeabilization Wash Buffer (BioLegend, USA) were used for intracellular staining following manufacturer’s recommendations. Cells were stained with anti-IFNγ-FITC (clone B27, BD Biosciences, USA), anti-TNFα-PE/Cy7 (clone Mab11, Thermo Fisher Scientific, USA), anti-GM-CSF-PE (clone BVD2-21C11, BioLegend, USA), anti-IL-2-PE/Cy7 (clone MQ1-17H12, BioLegend, USA).

### In vivo tumor model

NSG mice were injected with 2×10^6^ K562-A2/LMP2-GFP-luc2 tumor cells subcutaneously (s.c.). Eight days later, mice received 2×10^6^ human T cells (CD8:CD4 ratio = 3:1) intravenously (i.v.) in the tail vain. The mice were monitored for tumor growth by bioluminescent measurements (IVIS Lumina LT, Perkin Elmer, USA). Luciferin (Cambridge Bioscience, UK) was injected s.c. following manufacturer’s instructions. When tumor size reached 15mm in diameter measured with calipers, mice were sacrificed, and tumor tissue was isolated. Tumor growth rate was calculated by fitting an exponential function (y = c * e^bx^) in Microsoft Excel. The slope value corresponds to *b* in the exponential formula and was used as the tumor growth rate.

### Statistics

Prism version 8 software for Mac OS Catalina (GraphPad, USA) was used for two-way ANOVA with Tukey’s post-test for multiple comparisons. Flow cytometry data was analyzed by FlowJo 10 software (BD Biosciences, USA). Pie charts were created using Pestle and SPICE 6 software for Mac OS Catalina [81]. Duplicate measurements from one representative donor are shown in figures 3 and 4. Two other donors are shown in the supplement figures S2 and S3.

### Study approval

Protocols using live animals were performed in accordance with a United Kingdom Home Office project licence (PPL – P38881828) and the regulations of the University of Edinburgh Animal Welfare and Ethics Review Board, which approved these experiments.

## Authors contributions

SP designed research study, conducted experiments, acquired and analyzed data, generated figures, and wrote the manuscript. NL, DW, ART helped performing experiments. HS provided the TCR constructs and tumor cells. RZ designed research study, analyzed data and wrote the manuscript. All authors read and discussed manuscript drafts and approved the final version of the manuscript.

## Acknowledgments

The authors thank all their colleagues for helpful discussion, particularly current and former members of the Zamoyska lab. This work was supported by a project grant from Cancer Research UK (C25969/A23235, awarded to RZ) and a Wellcome Trust Investigator Award (WT205014/Z/16/Z, awarded to RZ).

## Conflict of interest

The authors have declared that no conflict of interest exists.

---

**Supplement Figure 1.**
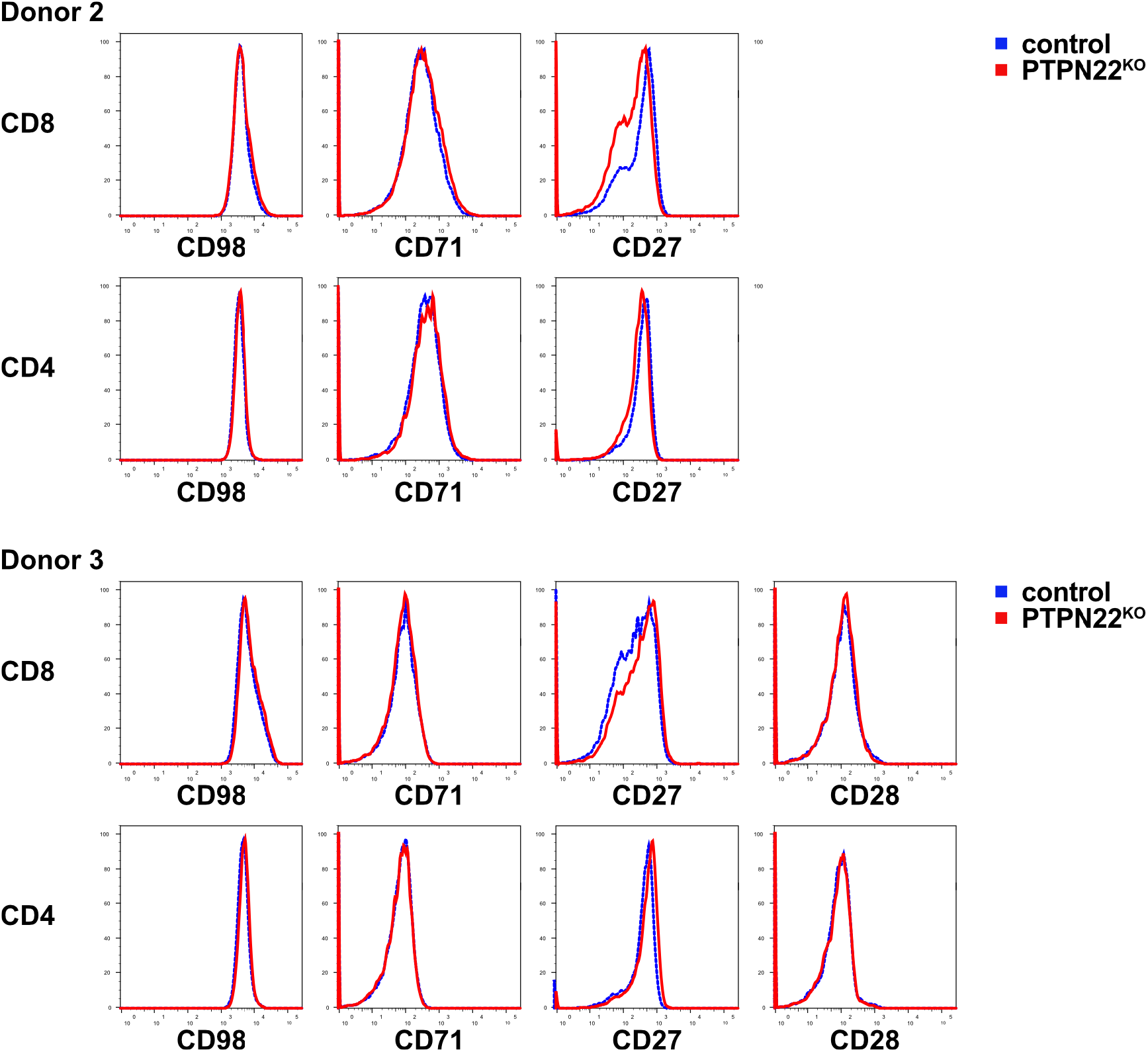
Surface marker expression on PTPN22^KO^ and control T cells. Expression of CD98, CD71, CD27, and CD28 was evaluated by cell surface staining and flow cytometry. Data of 2 different healthy donors are shown.

**Supplement Figure 2.**
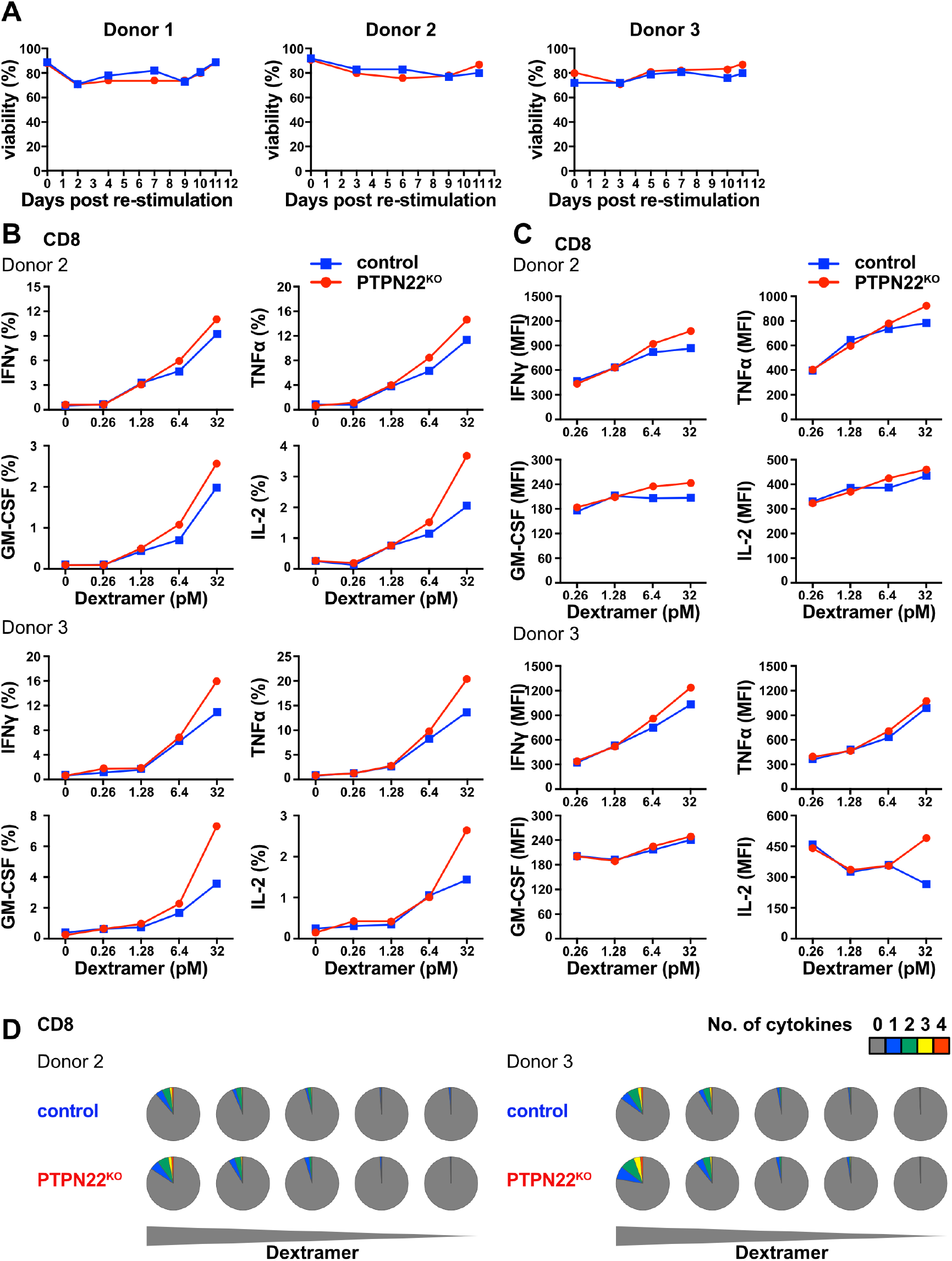
(A) Viability of PTPN22^KO^ and control T cells. Viability for 3 different donors was determined during *in vitro* expansion. PTPN22^KO^ and control T cells were re-stimulated for 4 hours with increasing concentrations of specific Dextramers. Frequencies of IFN^+^, TNF^+^, GM-CSF^+^ and IL-2^+^ T cells (B) and the mean fluorescence intensities (MFIs) of the cytokine-positive cells (C) were evaluated by intracellular staining and flow cytometry. Data are shown from duplicate measurements of two independent healthy donors. (D) Polyfunctional cytokine expression for CD8 T cells was calculated using FlowJo software and visualized with Pestle and SPICE 6 software. Pie chart fragments represent the fraction of cells expressing the number of cytokines indicated in the key. Data are shown from duplicate measurements of two independent healthy donors.

**Supplement figure 3.**
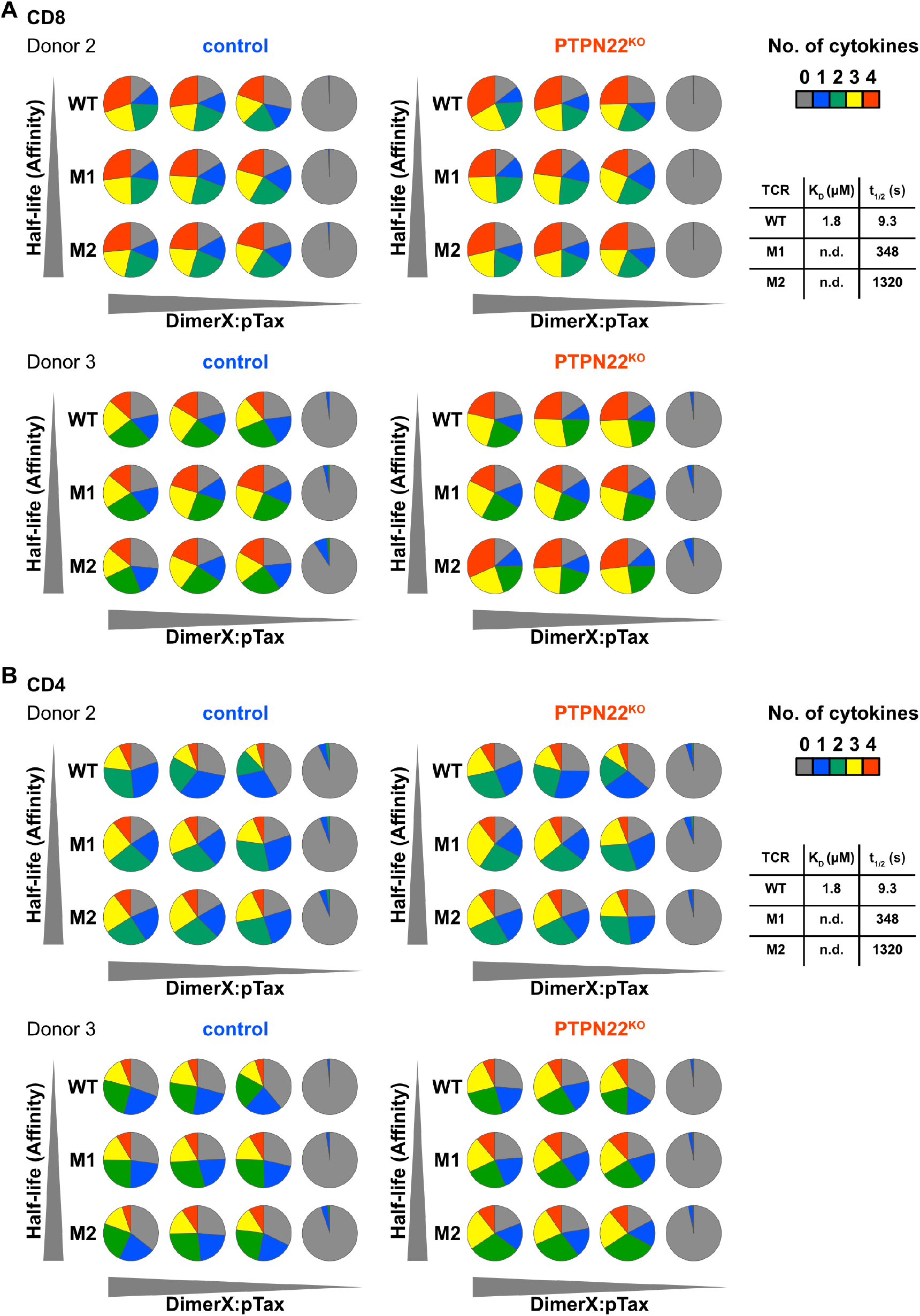
PTPN22^KO^ and control T cells were transduced with Tax-TCRs that have increasing affinities for the Tax antigen (WT<M1<M3) and re-stimulated with increasing concentrations of DimerX+pTax for 4 hours. Polyfunctional cytokine expression for CD8 (A) and CD4 (B) T cells were calculated using FlowJo software and visualized with Pestle and SPICE 6 software. Pie chart fragments represent the fraction of cells expressing the number of cytokines indicated in the key. Data are shown from duplicate measurements of two independent healthy donors.

## Notes

### Competing Interest Statement

The authors have declared no competing interest.

